# Labeling subcellular structures in living specimens using live-cell incompatible dyes with excellent optical properties

**DOI:** 10.1101/2020.05.10.086538

**Authors:** Yubing Han, Zhimin Zhang, Wenjie Liu, Yuanfa Yao, Yuchen Chen, Xin Luo, Wensheng Wang, Yingke Xu, Xu Liu, Cuifang Kuang, Xiang Hao

**Affiliations:** State Key Laboratory of Modern Optical Instrumentation, College of Optical Science and Engineering, Zhejiang University, Hangzhou, Zhejiang 310027, China; Department of Biomedical Engineering, Key Laboratory of Biomedical Engineering of Ministry of Education, Zhejiang Provincial Key Laboratory of Cardio-Cerebral Vascular Detection Technology and Medicinal Effectiveness Appraisal, Zhejiang University, Hangzhou, Zhejiang 310027, China; Ningbo Research Institute, Zhejiang University, Ningbo 315100, China; Collaborative Innovation Center of Extreme Optics, Shanxi University, Taiyuan 030006, China

## Abstract

Despite the urgent needs of imaging living specimens for cutting-edge biological research, most of the existing fluorescent labeling methods suffer from either poor optical properties or complicated operations to realize cell-permeability and specificity. Here, we introduce a method to overcome this tradeoff by incubating living cells and tissues with fluorescent dyes, no matter if they are cell-permeable or not, at particular conditions (concentration and temperature) without physical cell-penetration or chemical modifications. Based on this method, the mitochondrial labeling capability of Atto dyes, especially Atto 647N with extraordinary optical properties, together with interesting interactions between organelles is revealed. These results indicate the great potential of using dyes, which are normally considered “live-cell incompatible”, to capture the morphology and dynamics of subcellular structures in living specimens.

## Introduction

Biologists rely on an array of fluorescent microscopy to observe morphologies and dynamics in living cells, which is crucial in interpreting vital physiological and pathological activities. However, although dramatic improvements have been implemented since the seminal discovery of fluorophores and their applications in microscopy (*1,2*), it is still quite challenging to achieve live-cell specific staining simultaneously with high brightness and photostability.

In fact, most live-cell labeling methods, including fluorescent proteins (*3*), chemical tag techniques using cell-permeable fluorescent dyes (e.g., SNAP-Cell 647-SiR (*4*)), and live-cell organic fluorescent probes (e.g., MitoTracker dyes (*5*)), suffer from relatively low brightness and photostability (*6,7*). Poor optical properties of the fluorescent probes may lead to low signal-to-noise ratios (SNR) and fast photo-bleaching. Lower SNR reduces the quality of the microscope images or even introduce artifacts. Worse still, the photobleaching is undesirable in the long-term experiments. A dosage increase can partially remedy this problem, but it may also lead to non-specific labeling (*7,8*). Another option is to increase the illumination laser power, but it, in turn, further accelerates the photobleaching and introduces stronger phototoxicity. This issue stands out especially in the case of imaging methods for more information in multiple dimensions (e.g., high spatiotemporal resolution, long acquisition times, and large volume imaging) (*6,9,10*).

In the past few decades, various types of commercially available dyes, with much higher brightness and photostability than most of the live-cell probes, have been developed and widely used in fixed cells and organisms (*11*). However, most of these fluorophores were supposed to be “live-cell compatible”, and their intracellular cytosolic delivery and specifically labeling rely heavily on specific physical or chemical methods, such as microinjection, encapsulating vesicles, or chemical modifications using cell-penetrating peptides (*7,12,13*).

To resolve this dilemma, we present a novel strategy to label various types of subcellular structures both in living cells and in tissues using the bright and photostable dyes, which normally require complex delivery methods for subcellular labeling. Specifically, by incubating with living specimens at 1.5-15 μM for 30 min at 37°C or 20°C, the “live-cell incompatible” dyes can also be live-cell compatible. With multiple applications using our labeling strategy, these dyes (e.g., Atto 647N) shows great potential as a live-cell mitochondrial marker with excellent optical properties.

## Results

### Evaluation of different dyes for labeling living cells

Several types of fluorescent dyes were characterized according to their structure series.

Many cell-permeable cationic dyes (e.g., Rhodamine and Carbocyanine derivates) have been developed as mitochondrial probes, as they tend to accumulate in the mitochondrial matrix driven by the potential gradient in mitochondria at about 100-nM concentrations (*5,14,15*). Endoplasmic reticulum (ER) staining can also be realized by raising the dosages of these dyes (*7*). However, for the “live-cell compatible” dyes, concentrations at the 100-nM level are not enough.

Following our previous work (*7*), herein, aliquots of the indicated dyes were dissolved in Dimethyl sulfoxide (DMSO) to make 3-mM stock solutions, and these solutions were diluted with phenol red-free medium to 1.5-15 μM concentrations (DMSO/incubation buffer = 1/2,000) before use. After incubation with living U2OS (human osteosarcoma cell line) cells for 30 min at a proper temperature, a solution of Trypan blue (TB; 1 mg·mL^−1^ in PBS) was added to quench the fluorescence from the probes outside the cells (*7,16,17*).

Firstly, four Rhodamine derivatives, i.e., Atto 495, Atto 565, Atto 590, and Atto 647N N-Hydroxysuccinimide (NHS) esters, were investigated due to their relatively good optical properties in the corresponding spectral bands (Fig. 1 and figs. S1-6). When incubated at 37°C, which is the optimum temperature for cell culture, both line-shaped and bright dot signals inside the cells were observed (fig. S1B); whereas when the incubation temperature was set to 20°C (room temperature), the dot signals were dramatically reduced (fig. S1B). Mitochondrial staining was confirmed by colocalization experiments (Pearson correlation coefficient: 0.70, 0.57, and 0.82 for Atto 495, Atto 590, and Atto 647N, respectively; Fig. 1A-C and fig. S2), indicating their direct penetration through the plasma membrane. Besides, the dot signals were partially colocalized with endocytic-unassociated vesicles (early endosomes, late endosomes, and lysosomes; Figs. 1D-I and fig. S3-5), while no colocalization was found with other vesicular structures (fig. S6). Since the extracellular fluorescence (e.g. dye aggregates) was quenched by TB, these results indicate that (i) the dyes revealed by dot signals mainly entered living cells by endocytosis; (ii) incubation at relatively low temperature (e.g., 20°C) can effectively reduce endocytosis of these dyes so that they can mainly enter living cells through free diffusion based on the concentration gradient.

**Fig. 1.**
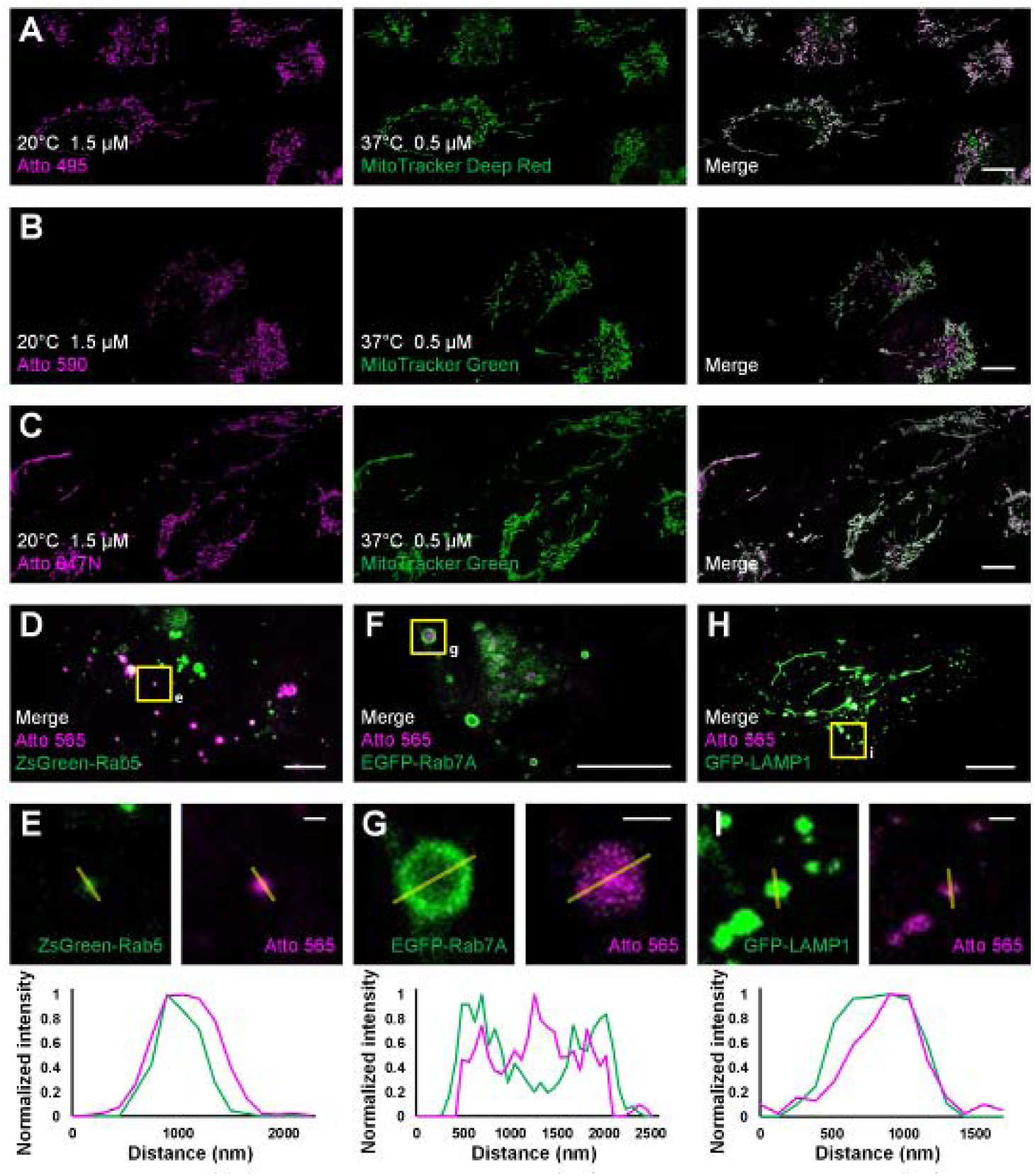
Co-localization studies employing different fluorescent probes and proteins as the standard mitochondrial markers. (**A**) Living U2OS cells were incubated with Atto 495 (magenta, 1.5 μM) for 30 min at 20°C and then with MitoTracker Deep Red (green, 0.5 μM) for 30 min at 37°C before imaging. Living U2OS cells were incubated with (**B**) Atto 590 or (**C**) Atto 647N (magenta, 1.5 μM) for 30 min at 20°C and then with MitoTracker Green (green, 0.5 μM) for 30 min at 37°C before imaging. Living U2OS cells transiently transfected by (**D**) ZsGreen-Rab5 (green; early endosomes), (**F**) EGFP-Rab7A (green; late endosomes), and (**H**) GFP-LAMP1 (green; lysosomes) were stained with Atto 565 (magenta, 6 µM) for 30 min at 37°C and imaged by confocal microscope. (**E, G, I**) Enlarged images from the boxed regions shown in d, f, h, respectively. Profiles represent the fluorescence intensities across the yellow lines in green and magenta channels. Scale bars, (**A**-**C, D, F, H**) 10 μm, and (**E, G, I**) 1 μm.

Then, as Carbocyanine dyes, anionic Alexa Fluor (AF) 647, zwitterionic Cy3B and cationic Cy5 were chosen as representative research objects (fig. S7A-C). No mitochondrial staining was found after the incubation at either 37°C or 20°C. For AF 647 and Cy3B, the introduction of sulfonic acid groups improves fluorescence and solubility in water but adds negative charges, preventing their affinity for mitochondria (*18*). For Cy5, the especially high hydrophobicity leads to its strong signals in the plasma membrane (fig. S7B), preventing its permeation into living cells (*19*).

Unexpectedly, when incubated at the same concentrations as that for the above dyes, zwitterionic BODIPY 650/665 labeled not only mitochondria but also the ER (fig. S7D), suggesting its higher cell-permeability than the other dyes tested. Our results indicate the potential of the BODIPY dyes as live-cell mitochondrial (or ER) markers, breaking through the early conclusion (*14*) that this purpose can only be achieved using cationic dyes.

In view of these observations, we hypothesize that without additional vector reagents and physical penetration, a group of fluorescent dyes, no matter if they were initially considered to be live-cell compatible or not, can mark subcellular structures in living cells using our labeling strategy (Table 1).

**Table 1.** Recommended conditions for specifically labeling subcellular structures in living cells using the live-cell incompatible dyes.

| *Subcellular structures* | *Electric charges* | *Additional properties* | *Incubation concentration* | *Incubation temperature* | *Recommended dyes* |
| --- | --- | --- | --- | --- | --- |
| *Mitochondria* | Cationic dyes, or zwitterionic BODIPY dyes | Moderate cell permeability | 1.5-15 μM | 20°C | Atto 647N |
| *ER* | Cationic dyes, or zwitterionic BODIPY dyes | Higher cell permeabilities or dosages | 15 μM | 20°C | BODIPY 650/665 |
| *Endocytic vesicles* | - | Particularly low cell-permeability | 1.5-15 μM | 37°C | AF 647, Cy3B |
| *Plasma membrane* | Cationic dyes | Especially high lipophilicity | 1.5-15 μM | 20°C | Cy5 |

### Labeling in various types of living cells

We also used the living HeLa (human adenocarcinoma cell line), HEK293 (human embryonic kidney epithelial cell line), NIH/3T3 (mouse embryonic fibroblast cell line) cells and primary cultural astrocytes from 1-day-old Sprague-Dawley (SD) rat (Fig. 2 and fig. S8). Notably, while the transfection in primary cultural astrocytes is challenging to achieve, the incubation with fluorescent probes is quite simple and efficient. Mitochondrial hyperfusion, which occurs upon cellular stresses and confers stress resistance (*20*), in astrocytes were captured (Fig. 2B-E and Movie S1). Some mitochondria moved radially (the direction is highlighted by the white arrowheads in Fig. 2D), and then either fused with other mitochondria (the blue arrowheads in Fig. 2D) or straighten out again (the yellow arrowheads in Fig. 2E). These dynamics may explain how mitochondrial hyperfusion is formed.

**Fig. 2.**
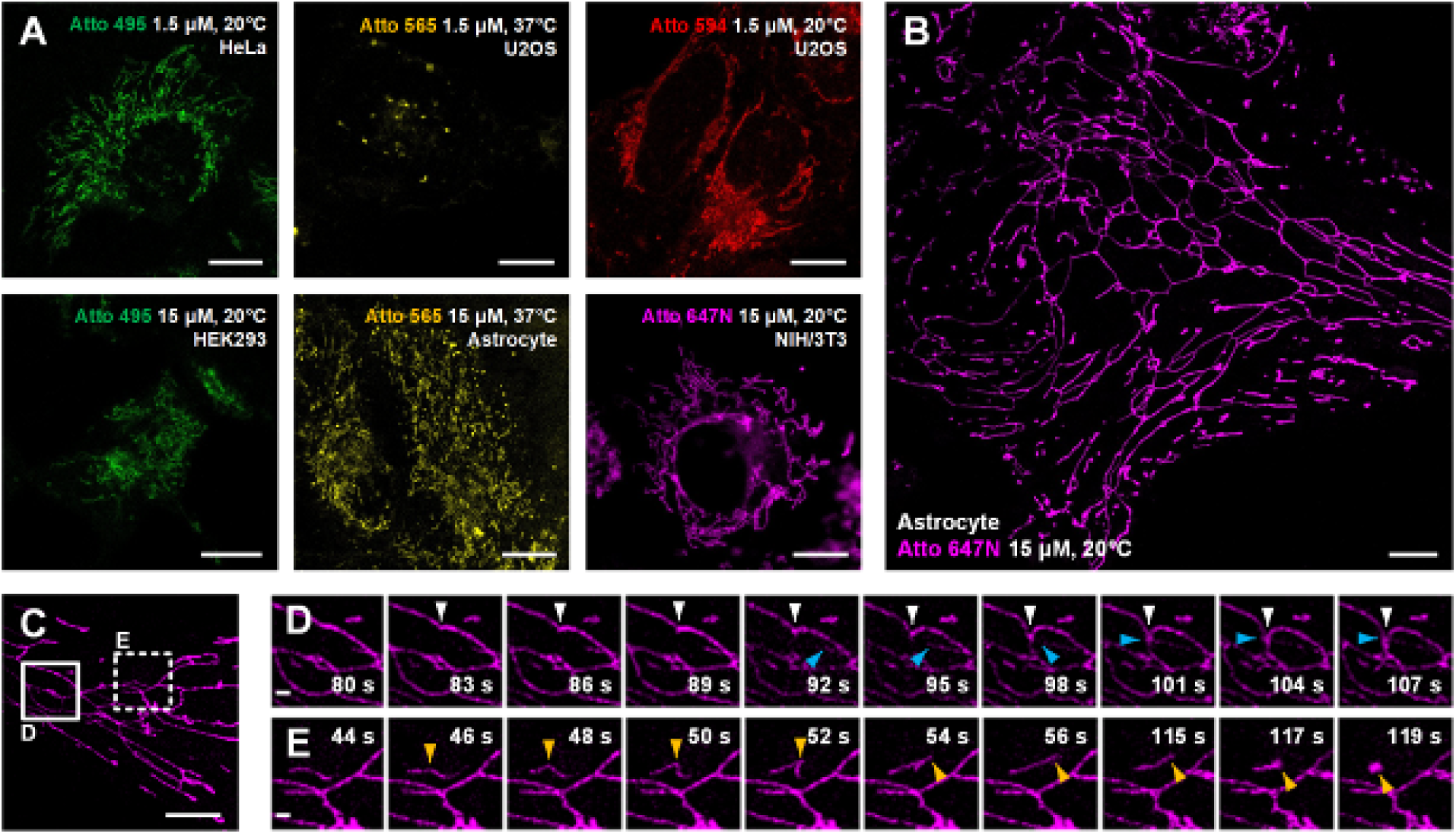
Various types of living cells stained with the Atto dyes. (**A**) Confocal images of living U2OS, Astrocyte, H3K293, HeLa, and NIH/3T3 cells incubated with the indicated Atto dyes at either 37°C or 20°C. (**B, C**) Living Astrocyte cells stained with Atto 647N at 20°C and imaged with a confocal microscope. (**D, E**) Enlarged time-lapse images from the solid and dashed boxed regions shown in **c**. No deconvolution or bleaching compensation procedure was applied during the figure rendering. For the time-lapse images, consecutive frames spaced at 1-s intervals were obtained; representative images of consecutive frames are displayed (more frames are shown in Movie S1). Scale bars, (**A-C**) 10 μm, and (**D, E**) 1 μm.

### Cell viability after labeling

Although raising the incubation concentrations helps the above dyes to permeate into living cells, it may also cause cytotoxicity from both the dyes themselves and the organic solvent in the incubation buffer (e.g., DMSO). To further confirm that our approach is compatible with the living specimens, the cytotoxicity of this method was then investigated by 3-(4,5-dimethylthiazol-2-yl)-5-(3-carboxymethoxyphenyl)-2-(4-sulfophenyl)-2H-tetrazolium (MTS) assay, which is a colorimetric method for determining the number of viable cells in proliferation or cytotoxicity assays (*21*). In living cells, the MTS tetrazolium compound can be converted into a colored formazan product that is soluble in the tissue culture medium, and the quantity of formazan product as measured by absorbance at 490 nm is directly proportional to the number of living cells in culture (*22*). After incubation with the indicated dyes at 37°C for 30 min (cells were untreated in the control groups), the cells were incubated with MTS reagents for another 3 h according to the instructions. The cell viabilities were expressed as the percentage of the absorbance of the dye-treated cells (after subtracting the values of the dyes in the cells) to the untreated controls, and all of the measurements were performed in triplicate. The results showed that after treated by the indicated dyes, the cell viabilities of the above five types of cells were usually higher than 80%, except in isolated cases (fig. S9). The cell viability higher than 100% (especially obvious for Atto 565 that enters the living cells mainly by endocytosis at 37°C) may result from the involvement of cellular redox factors in endocytosis (*23*). The result indicates the low cytotoxicity of our method.

### Optical properties of Atto 647N

We quantitatively compared the optical properties of the frequently-used red-absorbing fluorophores. In terms of brightness, Atto 647N exhibits much better performance than the SiR dye (*4*) and MitoTracker Deep Red (*5*) (Table S1). To evaluate photostability, fluorescence intensity curves were extracted from 20-min confocal imaging of U2OS cells (Fig. 3A and fig. S10), where all the imaging settings (laser power, size of field-of-view, imaging speed, etc.) were retained to control variables (Table S2). The results suggest that Atto 647N exhibited the best photostability (Fig. 3A and fig. S10A). Besides, by adding antioxidant reagent Prolong Live (*24*), all the bleaching speeds of these dyes decelerated to some extent. However, the difference between the dyes remained unchanged (Fig. 3A). The performance of Atto 647N was further investigated using stimulated emission depletion (STED) microscopy (Fig. 3B-E). The distance of mitochondrial outer membranes was resolved as 116 nm in STED mode, while the confocal image showed blurred structures without fine structures (Fig. 3C, D). In addition, structures of folded inner membranes, cristae, which were arranged in groups, and the voids between them were visible in the STED image (white arrows in Fig. 2E) (*7,25,26*). Therefore, these results indicate that applying our method in live-cell mitochondrial staining, especially in long-term or super-resolution imaging, Atto 647N can substitute SiR dye by offering better optical properties.

**Fig. 3.**
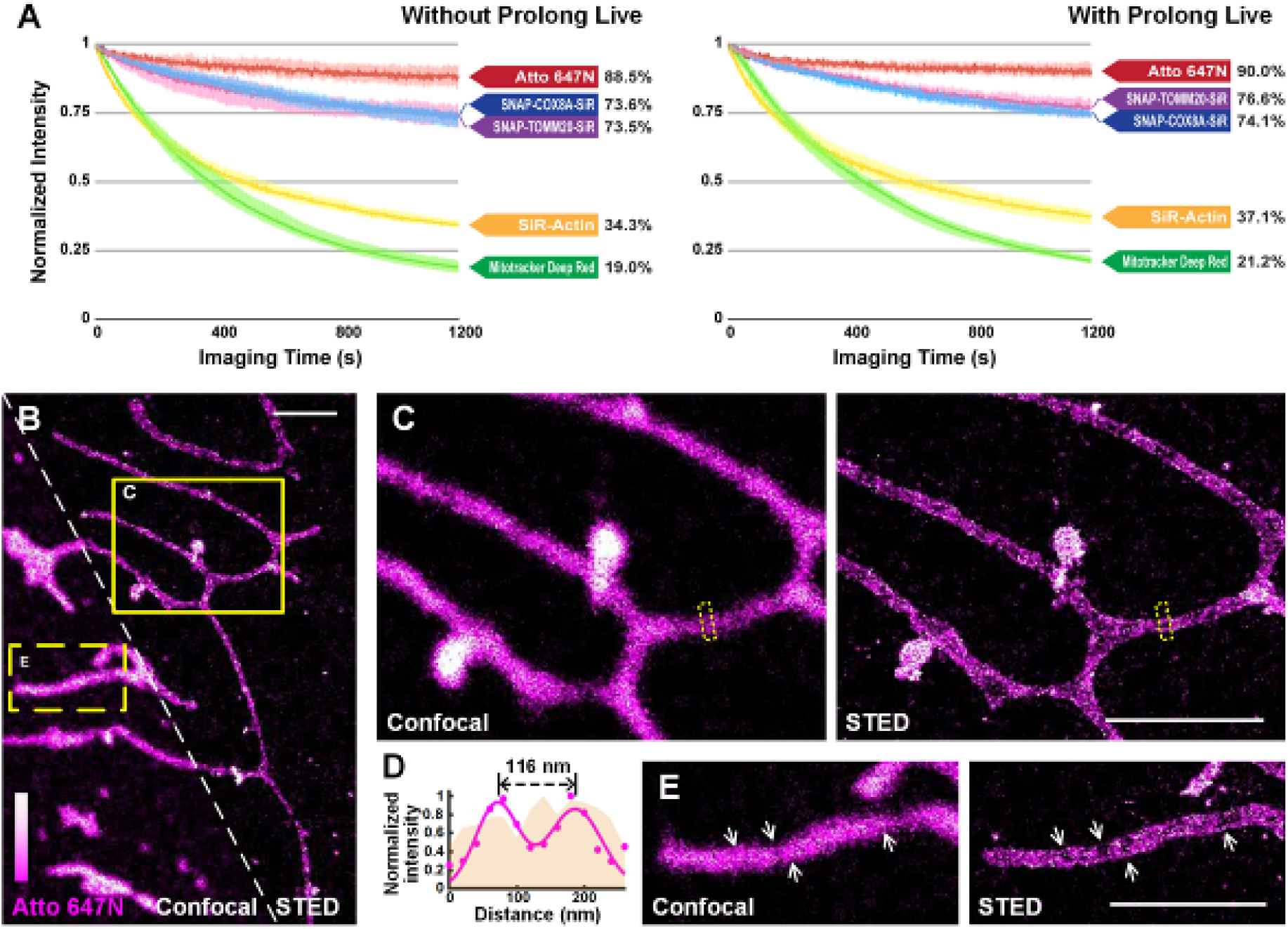
Atto 647N for live-cell imaging applications with high demand for optical properties. (**A**) Curves of intensity decay of the indicated dyes without (left) or with (right) the addition of Prolong Live under the same confocal imaging condition. Consecutive frames spaced at 1-s intervals were recorded. Error bars represent the standard deviations of triplicate experiments. (**B**) Living U2OS cells were incubated with Atto 647N for 30 min at 20°C. Enlarged confocal and STED images from the (**C**) solid or (**E**) dashed yellow boxes shown in b. (**D**) Intensity profiles at the positions within the dashed boxes in c (pink area for the confocal image, and magenta dots and Gaussian fitting curve for STED image). Scale bars, 2 μm.

### Dual-color applications

Dual-color staining combined with either fluorescent proteins (EMTB-3XGFP (*27*); Fig. 4A) or other probes (ER-Tracker Green; Fig. 4B-D and fig. S11) were performed, suggesting the compatibility of our method with other labeling strategies. Interestingly, the results in astrocytes indicate that some of the ER substructures colocalized with mitochondria over time (the white and yellow arrowheads in Fig. 4B and fig. S11A). The possibility of crosstalk between channels was ruled out since there were some places with no overlap of the two channels (highlighted with the yellow arrows in Fig. 4B). Besides, hitchhiking interactions (*27*) between ER-mitochondria (Fig 4C, D), ER-vesicles (white arrowheads in fig. S11C), and ER themselves (magenta arrowheads in fig. S11C) were observed in our experiments (Movies S2 and S3).

**Fig. 4.**
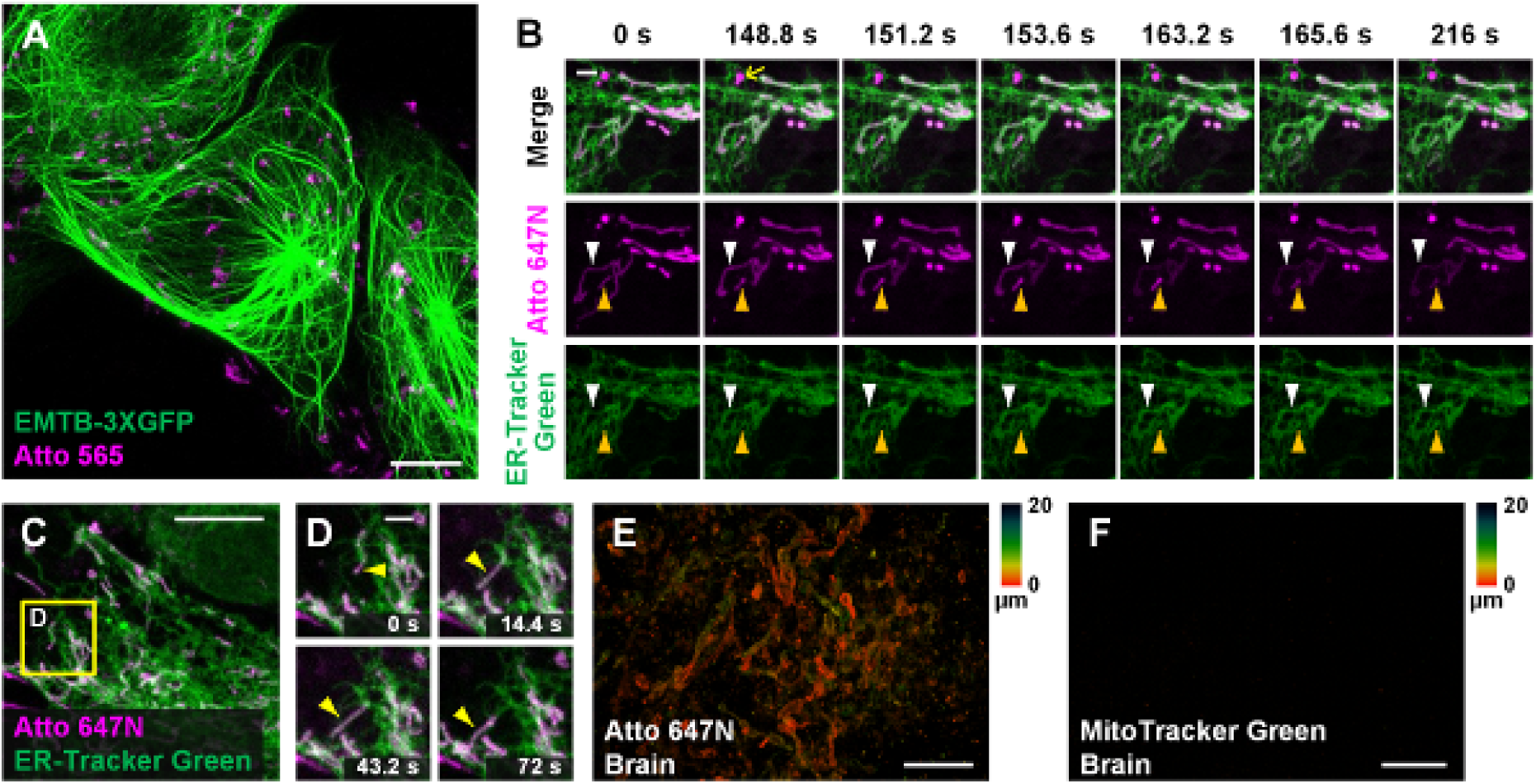
Other applications in living cells and tissues. (**A**) Dual-color confocal image (maximum intensity projection) of living U2OS cells labeled with EMTB-3XGFP (green) and Atto 565 (magenta; 3 μM). (**B, C**) Dual-color confocal image of living Astrocytes labeled with Atto 647N (magenta; 15 μM, middle row) and ER-Tracker Green (green; 2 μM, bottom row) at for 30 min at 20°C. (**D**) Mitochondria move along ER tubules. No deconvolution or bleaching compensation procedure was applied during the figure rendering. For the time-lapse images, consecutive frames spaced at 2.4-s intervals were recorded; representative images of consecutive frames are displayed (more frames are shown in Movies S2 and S3). Brain slices incubated with (**E**) Atto 647N and (**F**) MitoTracker Green. Color bars on the right of (**E**) and (**F**) indicate the imaging depth. Scale bars, (**A, C**) 10 μm, (**B, D**) 2 μm, and (**E, F**) 20 μm.

### Labeling living tissues

Compared to typical mammalian cells cultured on coverslips, labeling living tissue samples usually harbor a greater challenge. Tissues in vitro die faster than cultured cells without the support from the blood flow, resulting in faster mitochondrial potential gradient lost. As a consequence, the cationic dyes cannot affine to mitochondria. To permit sufficient staining deep into the living tissue, the fluorescent probes should permeate through the tissues fast enough so that they can bind to mitochondria before the gradient lost. Taking the brain slice from the mice for example, the brain slices were incubated with Atto 647N immediately after the mice were sacrificed. Our results suggest that 0.5-h incubation was sufficient enough for the dye to permeate for at least 20-μm thickness, indicating its excellent permeability through tissues (Fig. 4E and Movie S4). In contrast, MitoTracker Green failed to stain the brain sample (Fig. 4F), which suggests the abilities of Atto 647N to substitute MitoTracker probes in staining living tissues. Compared with the immunofluorescence techniques in previous works (*28*), our method requires fewer protocol steps, and the direct incubation used here avoids the structural change caused by fixation. Therefore, our method is more advantageous for many applications that expect instant actions, such as the rapid medical diagnoses of biospecimen in vitro and assessment for some mitochondrial diseases (*29,30*).

## Discussion

In summary, the capability of many fluorescent dyes, which was ignored to some extent before, to stain various types of subcellular structures in living specimens with high brightness and photostability is confirmed. Labeled structures include mitochondria, ER, endocytic vesicles, and the plasma membrane. The implementation requires only specific incubation conditions without any chemical modification or physical penetration, thereby minimizing the damages and artifacts induced during the sample preparation. Moreover, Atto 647N exhibited extraordinary brightness and photostability in live-cell mitochondrial labeling, which can substitute SiR dye in long-term imaging or super-resolution microscopy.

Due to the limits from objective conditions, especially that the structures of many of the commercially available fluorescent dyes are trade secret, not all of them are testified in this paper. However, the hypothesis built in this work gives a guideline to find these candidates out and to broaden the applications of existing dyes. Moreover, the phenomena observed here indicate their great potential to answer a wide range of biological questions in living cells and tissues.

## Materials and Methods

### Primary cultural astrocytes

Primary cultural astrocytes were obtained from Sprague-Dawley (SD) rat brains (1 day old). The rats were bought from Zhejiang Research Center of Laboratory Animals, China, sterilized with 75% ethanol and sacrificed by cutting off their heads. The following steps were all done on the ice. The scalps and skulls were incised, and the brains were taken out into pre-cooling Phosphate buffered saline (PBS; Thermo Fisher Scientific, Inc.). Then the cortex was isolated, cut into pieces using knives, and incubated with pre-warmed Trypsin-EDTA (Thermo Fisher Scientific, Inc.) at 37°C for 20 min. The mixture was centrifuged for 5 min at 1000 rpm, and the supernatant was removed. Then the cells were dissociated by adding 5 mL of pre-warmed growth medium (DMEM, high glucose + 10% fetal bovine serum (FBS); Thermo Fisher Scientific, Inc.) and vigorous pipetting. Finally, the cell suspension was incubated in a T25 flask (Thermo Fisher Scientific, Inc.) at 37°C in a humidified 5% CO_2_ environment. The medium was changed every two days, and after 7∼8 days, the culture flask was shaken manually for 30 min to remove the overlaying microglia exposed on the astrocyte layer. The supernatant containing microglia was discarded, and 5 mL of culture medium was added into the flask. This step was repeated for once to remove oligodendrocyte precursor cells.

### Cell culture

HeLa (human adenocarcinoma cell line), HEK293 (human embryonic kidney epithelial cell line), NIH/3T3 (mouse embryonic fibroblast cell line), and U2OS (human osteosarcoma cell line) cells were purchased from the American Type Culture Collection. HeLa, HEK293, and NIH/3T3 cells were cultured in the DMEM medium (Thermo Fisher Scientific, Inc.). U2OS cells were cultured in McCoy’s 5A medium (Thermo Fisher Scientific, Inc.). All media were supplemented with 10% (v/v) FBS, and the cultures were maintained at 37°C in a humidified 5% CO_2_ environment.

### Transfection

Cells were grown overnight in 24-well plates at 37°C in a 5% CO_2_ atmosphere. After reaching over 80% confluence, the plasmids mRuby-Clathrin (Addgene #55852), pEGFP-Sec23A (Addgene #66609), ZsGreen-Rab5 (custom synthesized by Genomeditech (Shanghai, China) Co., Ltd.), EGFP-Rab7A (Addgene #28047), GFP-LAMP1 (Addgene #16290), EMTB-3XGFP (Addgene #26741), pSNAPf-Cox8A (Addgene #101129), or pSNAPf-TOMM20 (custom synthesized by Genomeditech (Shanghai, China) Co., Ltd.) was transfected into the cells using Lipofectamine 3000 (Thermo Fisher Scientific, Inc.) according to the manufacturer’s instructions. After 24 h, the transfected cells were digested with trypsin-EDTA and seeded into Nunc Glass Bottom Dishes (Φ 12 mm, Thermo Fisher Scientific, Inc.) at a density of 1.5∼2.0 × 104 per well in growth medium (150 µL). The cells were grown for an additional 12∼24 h before incubation with the indicated probes.

### Live-cell labeling with commercially available probes

Before staining, the indicated cells were seeded in Nunc Glass Bottom Dishes at a density of 1.5∼2.0 × 10^4^ per well in growth medium (150 µL). After overnight incubation, the cells were washed three times with PBS. Work solutions of the indicated probes at different concentrations were prepared with phenol red-free DMEM (Thermo Fisher Scientific, Inc.). The cells were then incubated with MitoTracker Deep Red FM (200-500 nM, 100 µL; Thermo Fisher Scientific, Inc.), MitoTracker Green FM (1000 nM, 100 µL; Thermo Fisher Scientific, Inc.), SiR-actin (Cytoskeleton, Inc.) in a 5% CO_2_ atmosphere at 37°C for 30 min. For SNAP-Cell 647-SiR (New England Biolabs, Inc.) dye labeling, transfection of pSNAPf-Cox8A or pSNAPf-TOMM20 was performed before incubation of SNAP-Cell 647-SiR (3 µM) at 37°C for 30 min. After incubation, the supernatant was discarded, and a solution of Trypan blue (TB, 100 µL, 1 mg mL^-1^; Sigma-Aldrich Co., LLC) in PBS was added to exclude the dead cells and quench the extracellular fluorescence from the probes bound to either the cell membrane or the dish surface (7,16,17). After 1 min, TB was removed, and the cells were washed twice gently with PBS. Cells were immersed in Live Cell Imaging Solution (Thermo Fisher Scientific, Inc.) before imaging. ProLong Live Antifade Reagent (Thermo Fisher Scientific, Inc.) was added to this solution according to the manufacturer’s instructions when required.

### Live-cell labeling with fluorescent dyes

Before staining, the indicated cells were seeded in Nunc Glass Bottom Dishes at a density of 1.5∼2.0 × 10^4^ per well in growth medium (150 µL). After overnight incubation, the cells were washed three times with PBS. Aliquots of Atto 495, Atto 565, Atto 590, Atto 647N (Sigma-Aldrich Co., LLC), BODIPY 650/665, Alexa Fluor 647 (Thermo Fisher Scientific, Inc.), Cy3B, and Cy5 (GE Healthcare Co., Ltd) NHS esters were dissolved in Dimethyl sulfoxide (DMSO; Sigma-Aldrich Co., LLC) to make 3-mM stock solutions at −20°C. Stock solutions were diluted with phenol red-free DMEM to work solutions at different concentrations before use. The cells were incubated with these dyes either at 37°C in a 5% CO_2_ atmosphere or at 20°C for 30 min. The post-incubation treatment was the same as that of the commercially available probes above.

### Labeling living tissue slices with fluorescent probes

All animal procedures were conducted in compliance with the guidelines for animal care and use of Zhejiang University and conformed the Guide for the Care and Use of Laboratory Animal published by the National Academy Press (Washington, DC, 1996). The 12-week-old Institute of Cancer Research (ICR) mice were obtained from Zhejiang Academy of Medical Sciences [License Number: SCXK (Zhe) 2014001] and housed in cages under a standard condition of temperature (23 ± 2°C), relative humidity (55 ± 5%) and light 12/12 h light/dark cycle. The mice were free to access to food and water. Before the experiment, the mice were executed by euthanasia. The brains were separated and washed with cold PBS buffer. Then the brains were cut into slices from different orientations and incubated with Atto 647N (15 µM) for 30 min at 37°C. After incubation, the supernatant was discarded, and the tissue slices were washed twice gently with PBS. Then the tissue slices were put on the Nunc Glass Bottom Dishes with wound surface toward the glass bottom.

### MTS assay

The cytotoxicity of the Atto dyes on different cell lines was tested using the MTS assay (21). Cells (3-4 × 10^3^ cells per well) were seeded into a 96-well plate and cultured in growth medium for 24 h. The cells were incubated with the dyes (1.5 µM or 15 µM) in a 5% CO_2_ atmosphere at 37°C for 30 min, washed twice gently with PBS, and immersed in 100 µL of growth medium and 20 µL of CellTiter 96 AQueousOne Solution Reagent (Promega Co.). After another incubation for 3 h at 37°C in a 5% CO_2_ atmosphere, the absorbance was recorded at 492 nm using a TECAN GENios Plus ELISA reader (Tecan, Inc.). The cell viabilities were expressed as the percentage of the A492 of the dye-treated cells (after subtracting the values of the dyes, which was tested in cells without MTS treats) to the untreated controls, and all of the measurements were performed in triplicate.

### Confocal laser scanning microscopy

The confocal images were obtained using a C2 confocal laser scanning microscope (Nikon, Inc.) equipped with a 100×/1.49 numerical aperture oil immersion objective lens and were analyzed with NIS-elements (Nikon, Inc.) and ImageJ software (National Institutes of Health).

### STED microscopy

The STED images were obtained using a STEDYCON microscope (Abberior, GmbH.) equipped with a 100×/1.49 numerical aperture oil immersion objective lens and were analyzed with ImageJ software.

### Statistical Analysis

Statistical analysis was performed using Microsoft Excel 2019 (Microsoft Co., Ltd). Averages were represented as mean ± SD, and the number of replicates was indicated in respective figures and figure legends.

## Acknowledgments

We thank Mr. Yisheng Wu from SRstar Instruments Ltd., Shanghai, China, for operating the Abberior STEDYCON. This work was financially sponsored by the grants from National Key R&D Program of China (2018YFA0701400), the National Natural Science Foundation of China (61827825, 61735017, and 31901059), Fundamental Research Funds for the Central Universities (2019XZZX003-06 and 2019QNA5006), China Postdoctoral Science Foundation (2019M662042), Natural Science Foundation of Zhejiang province (LR16F050001), Zhejiang Lab (2018EB0ZX01), and ZJU-Sunny Photonics Innovation Center (2019-01).

## Author contributions

X. Hao, Y. Han, C. Kuang, and X. Liu conceived of the project. Experiments were performed primarily by Y. Han. Z. Zhang, W. Liu, Y. Chen, and W. Wang set up the system and contributed to the imaging. Y. Han, Y. Yao, and X. Luo prepared the samples. Y. Han and X. Hao drafted the manuscript. All authors contributed to the manuscript polish.

## Data and materials availability

The authors declare that all data supporting the findings of this study are available within the article and its Supplementary Information files or from the corresponding author X. H. on reasonable request.

