## Supplementary information for "Labeling subcellular structures in living specimens using live-cell incompatible dyes with excellent optical properties"

**Supplementary Figure 1**. Characterization of the Atto dyes in living cells........................................................... 2

**Supplementary Figure 2**. Co-localization studies employing MitoTracker dyes as the standard mitochondrial markers. ........................................................................................................................................ 3

**Supplementary Figure 3**. Co-localization studies employing ZsGreen-Rab5 as the standard early endosomal marker. ........................................................................................................................................ 4

**Supplementary Figure 4**. Co-localization studies employing EGFP-Rab7A as the standard late endosomal marker. ........................................................................................................................................ 5

**Supplementary Figure 5**. Co-localization studies employing GFP-LAMP1 as the standard lysosomal marker....... 6

**Supplementary Figure 6**. Co-localization studies employing mRuby-Clathrin and pEGFP-Sec23A as the standard markers for endocytic-unassociated vesicular structures........................................................................... 7

**Supplementary Figure 7**. Confocal images of living U2OS cells labeled with different dyes................................ 8

**Supplementary Figure 8**. Confocal images of different cell lines labeled with the Atto dyes. ................................ 9

**Supplementary Figure 9**. Cell viabilities of different cell lines stained with the Atto dyes. .................................. 10

**Supplementary Figure 10**. The first and last frames from 20-min confocal imaging of different probes.............. 13

**Supplementary Figure 11**. Dual-color confocal images of living Astrocytes......................................................... 14

**Supplementary Figure 12**. Applications in living brain slices. ................................................ **错误!未定义书签。**

**Supplementary Table 1**. Comparison of the optical properties of frequently-used red-absorbing fluorescent dyes. ........................................................................................................................................ 15

**Supplementary Table 2**. Experimental conditions for long-term confocal imaging............................................... 15

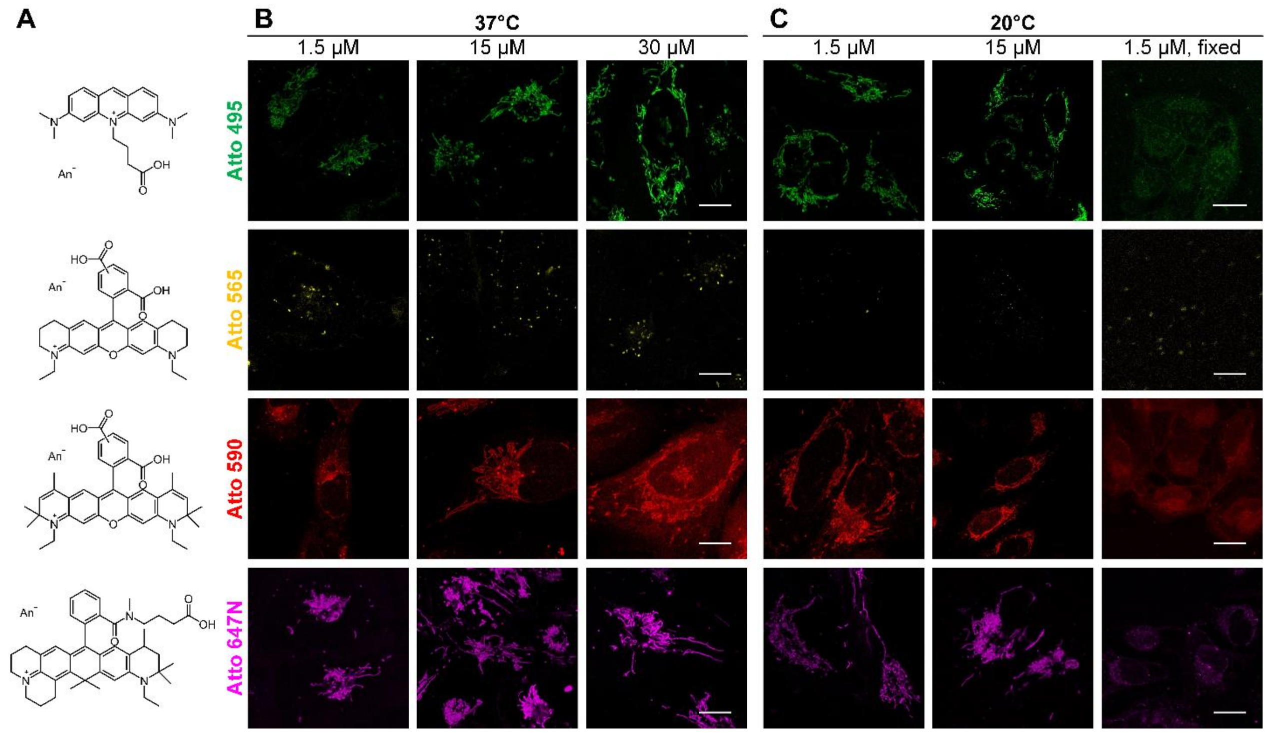


**Figure S1**. Characterization of the Atto dyes in living cells.

(**A**) Chemical structures of Atto 495, Atto 565, Atto 590, Atto 647N. Confocal images of living U2OS cells incubated with the Atto dyes at either (**B**) 37°C or (**C**) 20°C for 30 min. The fixation was accomplished by incubation with 4% paraformaldehyde for 10 min. Scale bars: 10 µm.

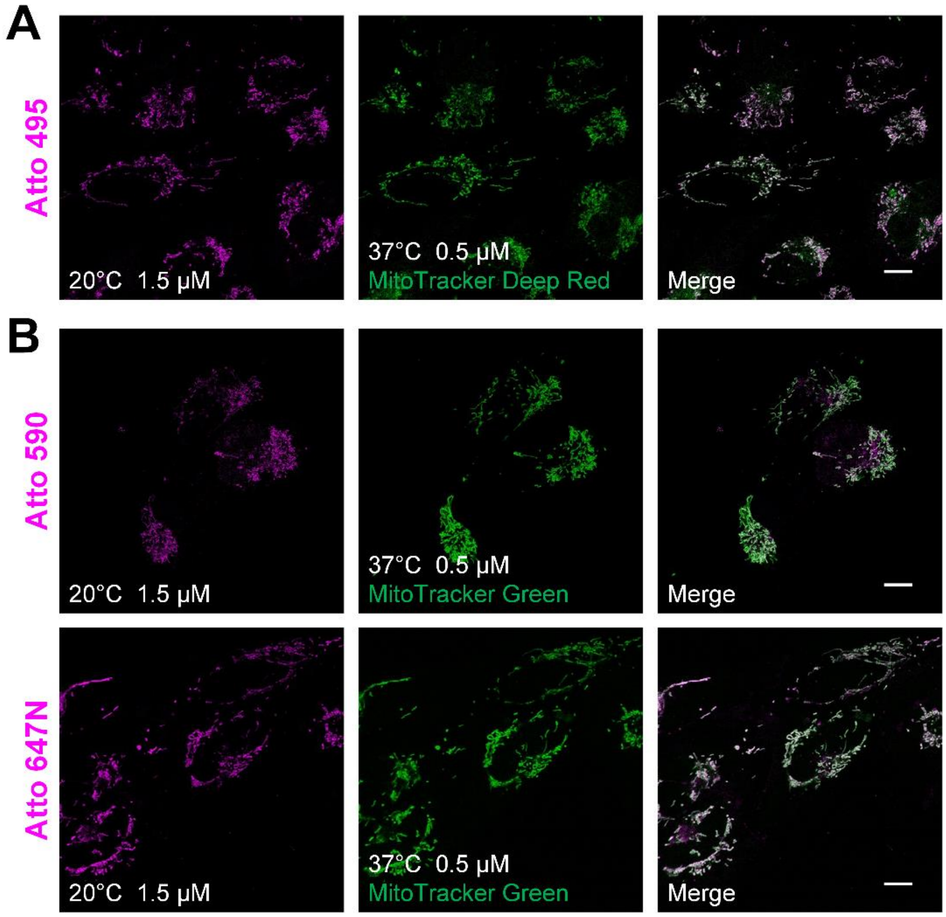


**Figure S2**. Co-localization studies employing MitoTracker dyes as the standard mitochondrial markers.

(**A**) Living U2OS cells were incubated with Atto 495 (magenta, 1.5 μM) for 30 min at 20°C and then with MitoTracker Deep Red (green, 0.5 μM) for 30 min at 37°C before imaging. (**B**) Living U2OS cells were incubated with Atto 590 or Atto 647N (magenta, 1.5 μM) for 30 min at 20°C and then with MitoTracker Green (green, 0.5 μM) for 30 min at 37°C before imaging. Scale bars: 10 μm.

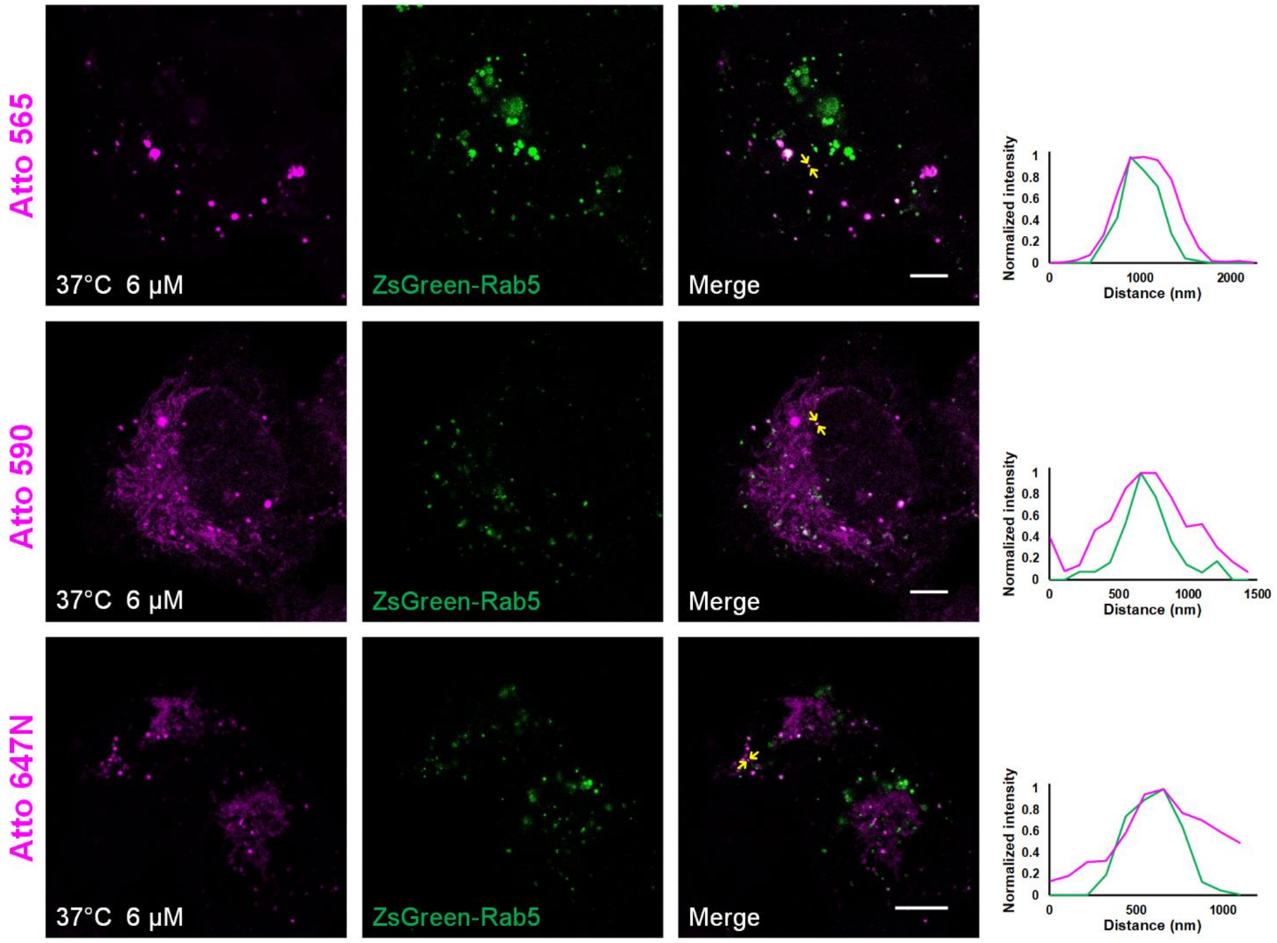


**Figure S3**. Co-localization studies employing ZsGreen-Rab5 as the standard early endosomal marker.

Living U2OS cells transiently transfected by ZsGreen-Rab5 (green, the second column from left) were stained with Atto 565, Atto 590, or Atto 647N (magenta, 6 µM, the first column) for 30 min at 37°C and imaged by confocal microscope. On the right, the intensity profiles at the position denoted by the yellow arrows in the merged image (the third column from left) are shown, indicating the co-localization of the Atto dyes and the early endosomes. Scale bars: 10 µm.

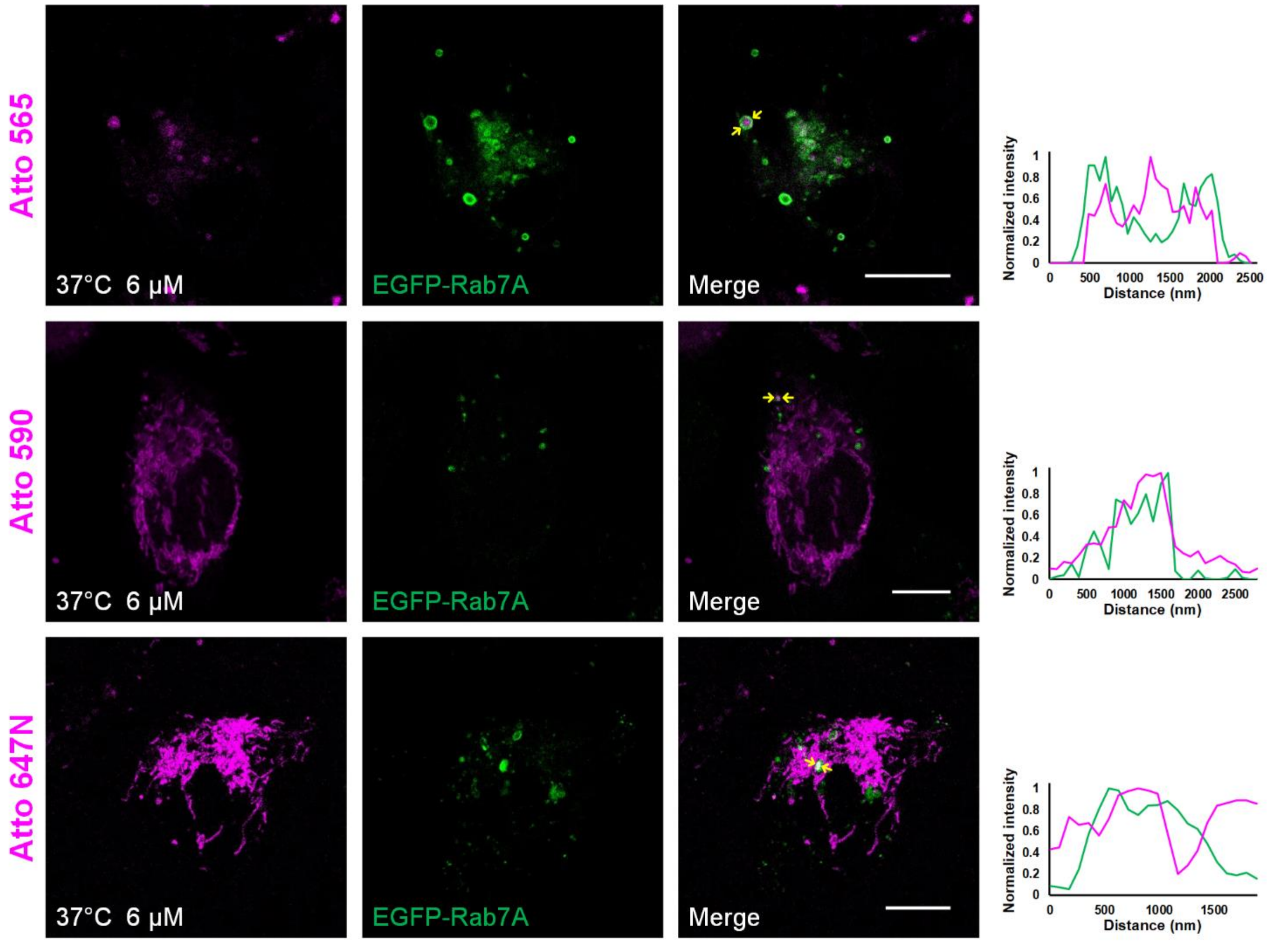


**Figure S4**. Co-localization studies employing EGFP-Rab7A as the standard late endosomal marker.

Living U2OS cells transiently transfected by EGFP-Rab7A (green, the second column from left) were stained with Atto 565, Atto 590, or Atto 647N (magenta, 6 µM, the first column) for 30 min at 37°C and imaged by confocal microscope. On the right, the intensity profiles at the position denoted by the yellow arrows in the merged image (the third column) are shown, indicating the co-localization of the Atto dyes and the late endosomes. Scale bars: 10 µm.

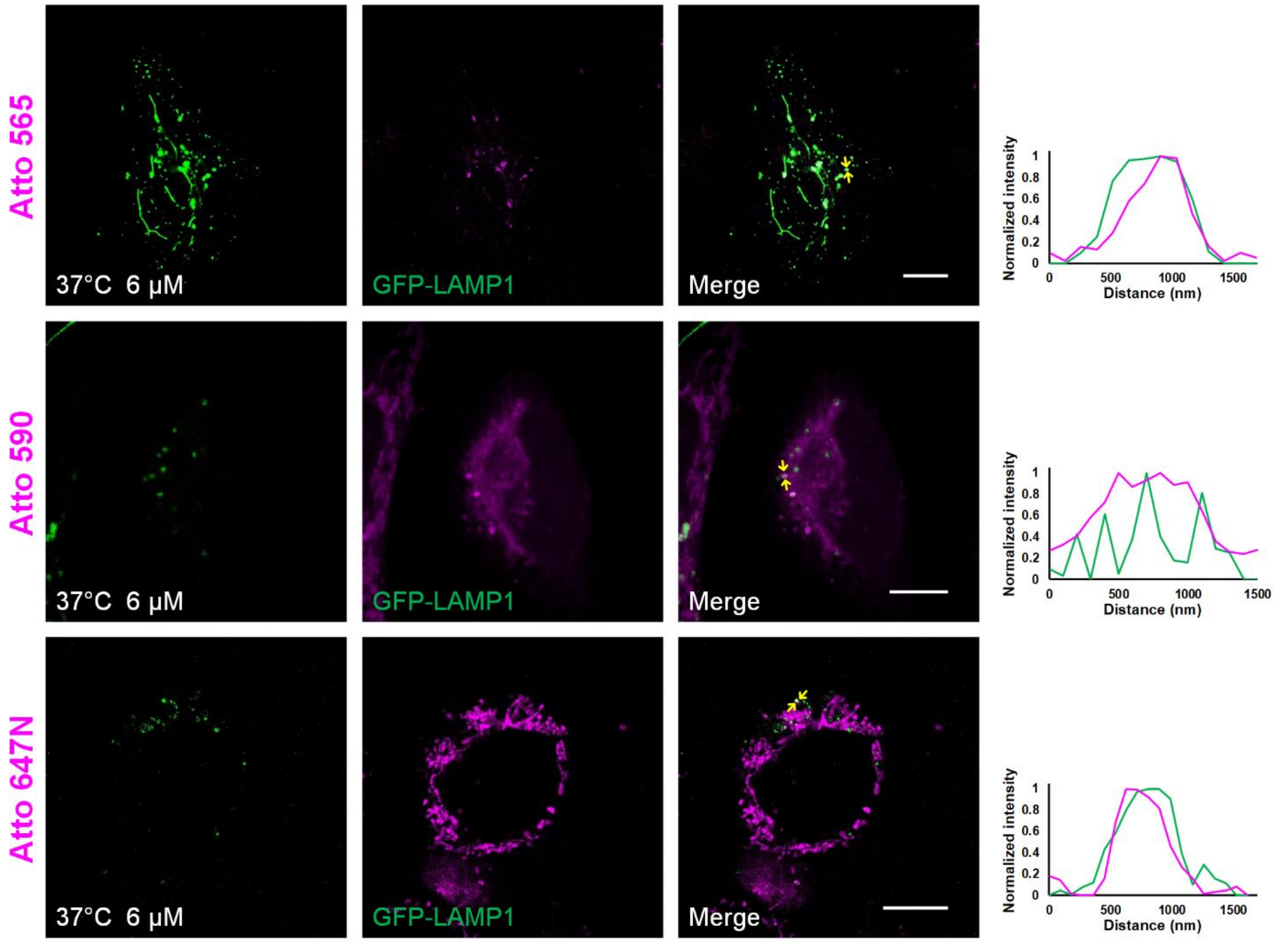


**Figure S5**. Co-localization studies employing GFP-LAMP1 as the standard lysosomal marker.

Living U2OS cells transiently transfected by GFP-LAMP1 (green, the second column from left) were stained with Atto 565, Atto 590, or Atto 647N (magenta, 6 μM, the first column) for 30 min at 37°C and imaged by confocal microscope. On the right, the intensity profiles at the position denoted by the yellow arrows in the merged image (the third column) are shown, indicating the co-localization of the Atto dyes and the lysosomes. Scale bars: 10 μm.

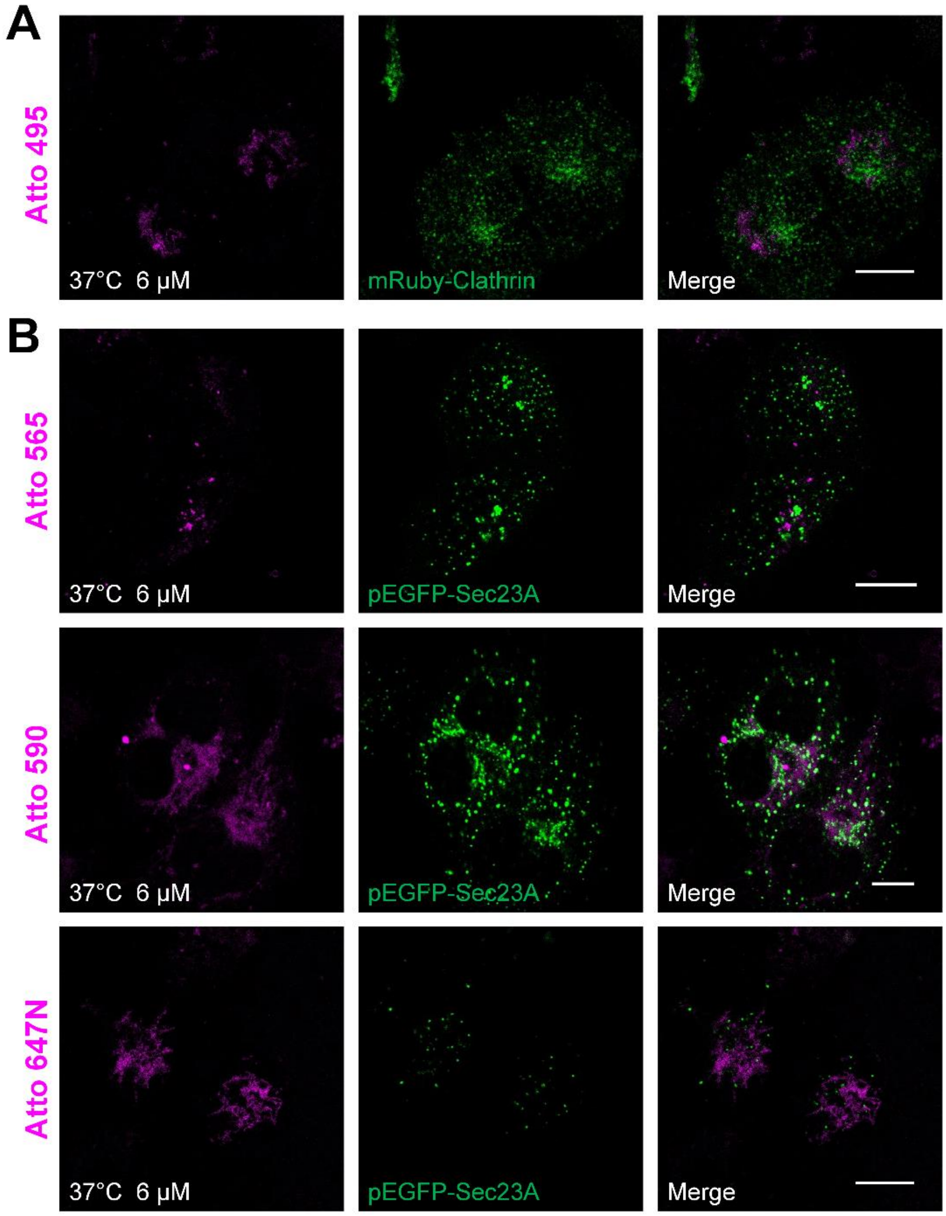


**Figure S6**. Co-localization studies employing mRuby-Clathrin and pEGFP-Sec23A as the standard markers for endocytic-unassociated vesicular structures.

(**A**) Living U2OS cells transiently transfected by mRuby-Clathrin (green, the middle sub-figure) were stained with Atto 495 (magenta, 6 μM, the left sub-figure) for 30 min at 37°C and imaged by confocal microscope. Two images were then merged and shown as the sub-figure on the right. Similarly, in (**B**), living U2OS cells transiently transfected by pEGFP-Sec23A (green, the middle column) were stained with Atto 565, Atto 590, or Atto 647N (magenta, 6 μM, the left column) for 30 min at 37°C and imaged by confocal microscope. The merged images were shown on the right in each row. Scale bars: 10 μm.

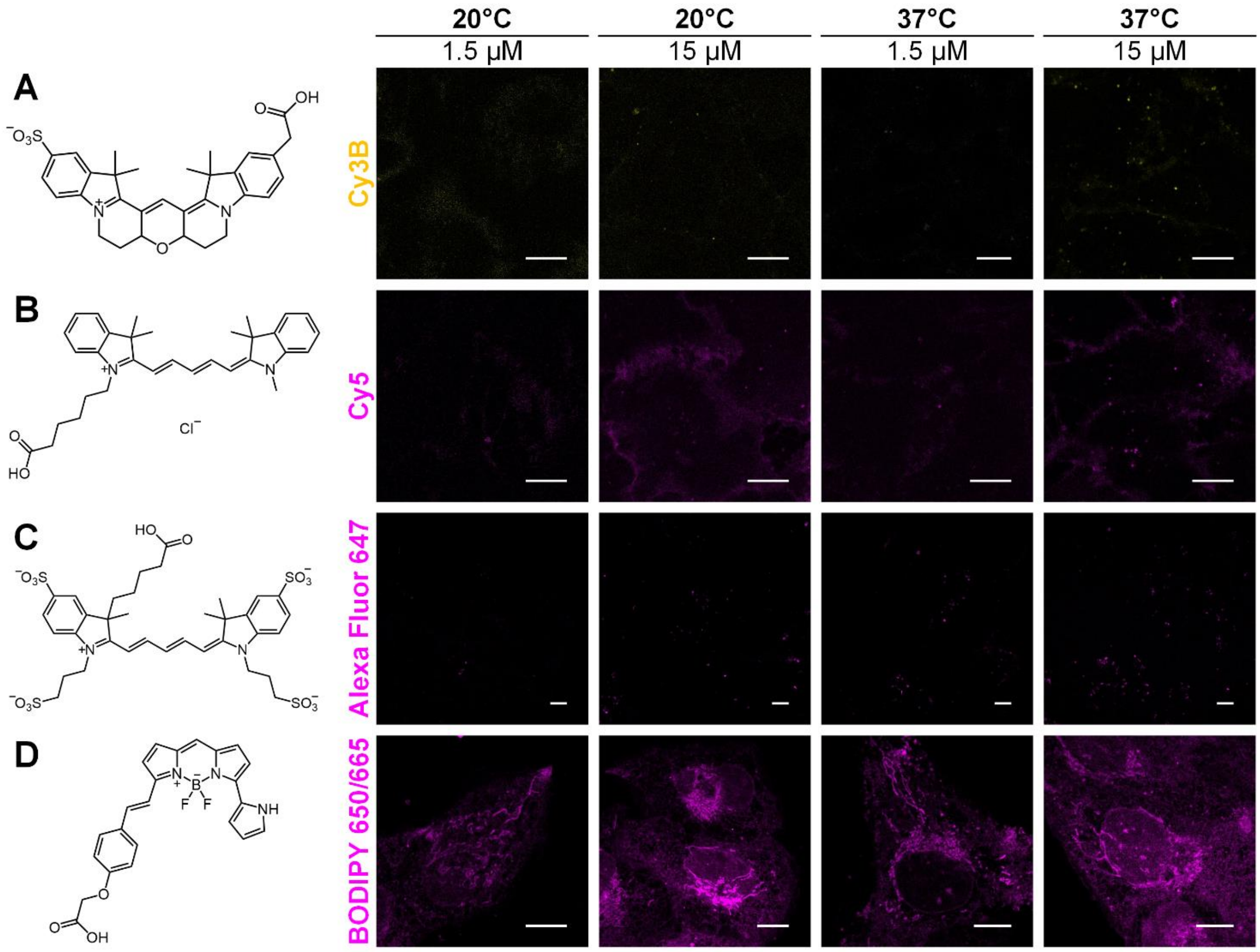


**Figure S7**. Confocal images of living U2OS cells labeled with different dyes.

Cells were incubated with (**A**) Cy3B, (**B**) Cy5, (**C**) Alexa Fluor 647, and (**D**) BODIPY 650/665 at different concentrations (1.5 µM or 15 µM) and temperatures (20°C or 37°C). Scale bars: 10 µm.

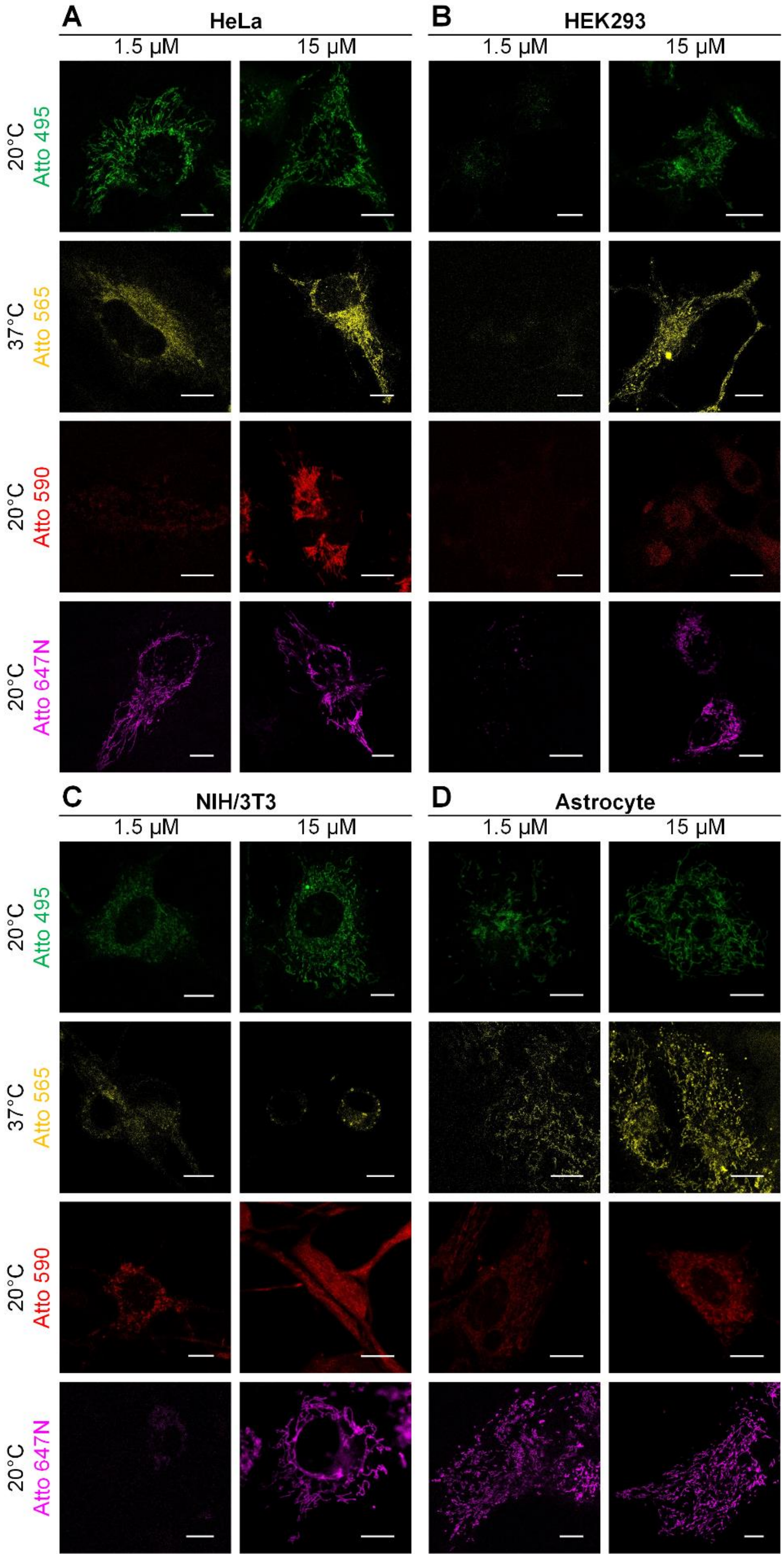


**Figure S8**. Confocal images of different cell lines labeled with the Atto dyes.

Living (**A**) HeLa, (**B**) HEK293, (**C**) NIH/3T3, or (**D**) Astrocyte cells were incubated with Atto 495 (at 20°C), Atto 565 (at 37°C), Atto 590 (at 20°C), and Atto 647N (at 20°C) for 30 min. Scale bars: 10 μm.

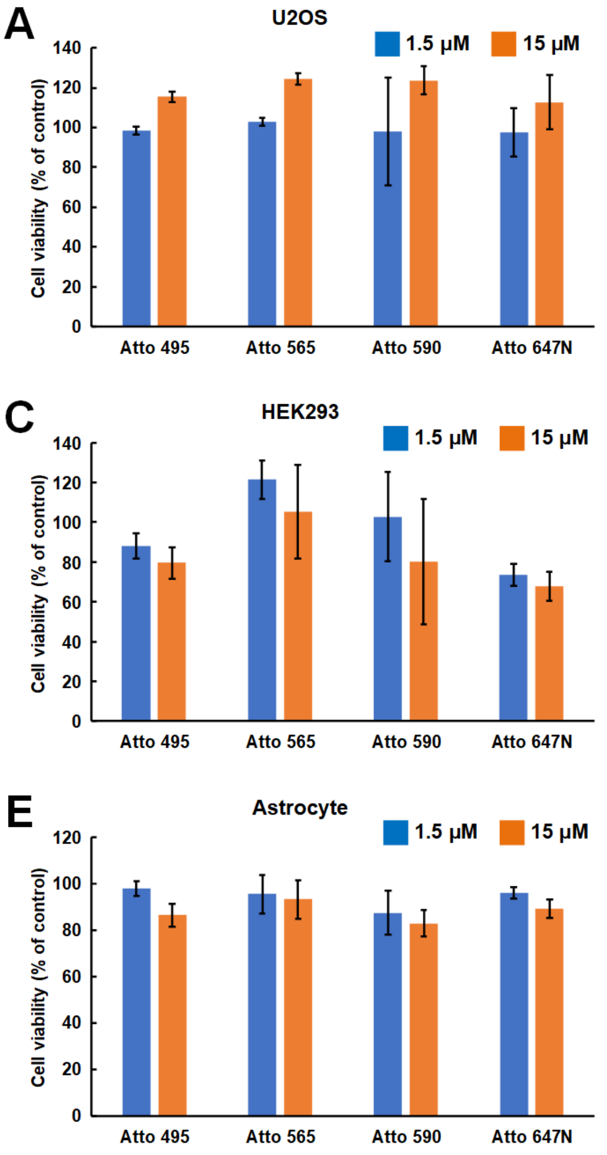


**Figure S9**. Cell viabilities of different cell lines stained with the Atto dyes.

Different living cells, (**A**) U2OS, (**B**) HeLa, (**C**) HEK293, (**D**) NIH/3T3, or (**E**) Astrocyte, were incubated with the Atto dyes at 37°C for 30 min. Error bars represent the standard deviations of triplicate experiments.

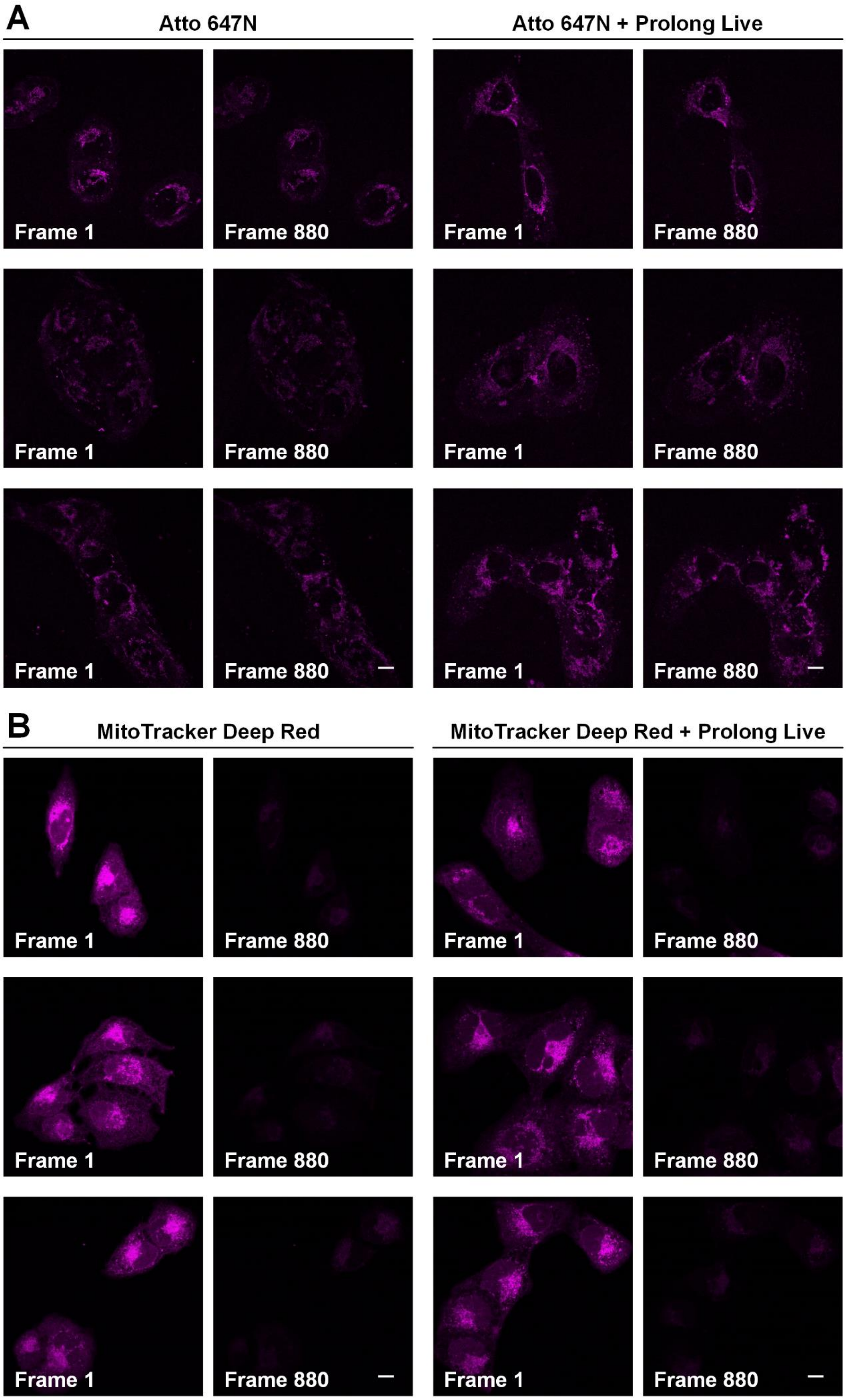
A
Atto 647N
Atto 647N + Prolong Live
Frame 1
Frame 880
B
MitoTracker Deep Red
MitoTracker Deep Red + Prolong Live

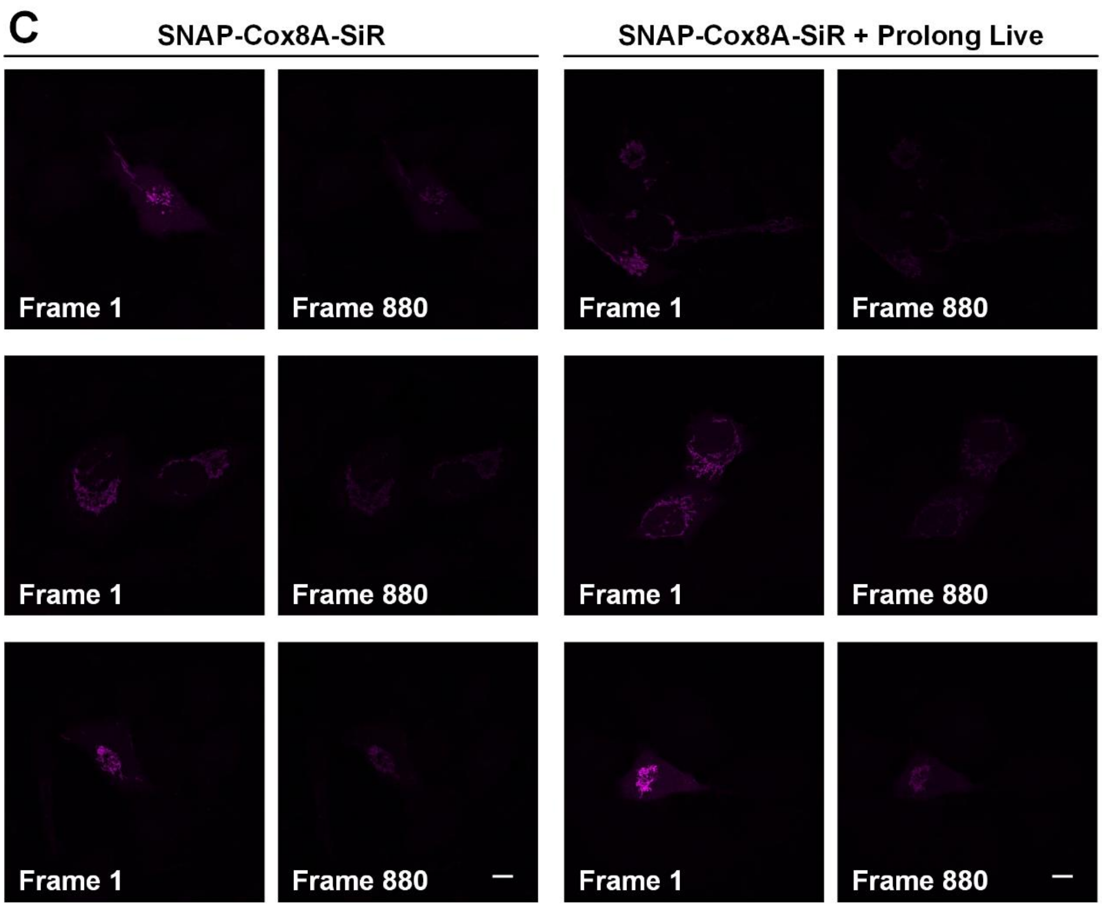
C
SNAP-Cox8A-SiR
SNAP-Cox8A-SiR + Prolong Live
Frame 1
Frame 880
Frame 1
Frame 880
Frame 1
Frame 880
Frame 1
Frame 880
Frame 1
Frame 880
Frame 1
Frame 880

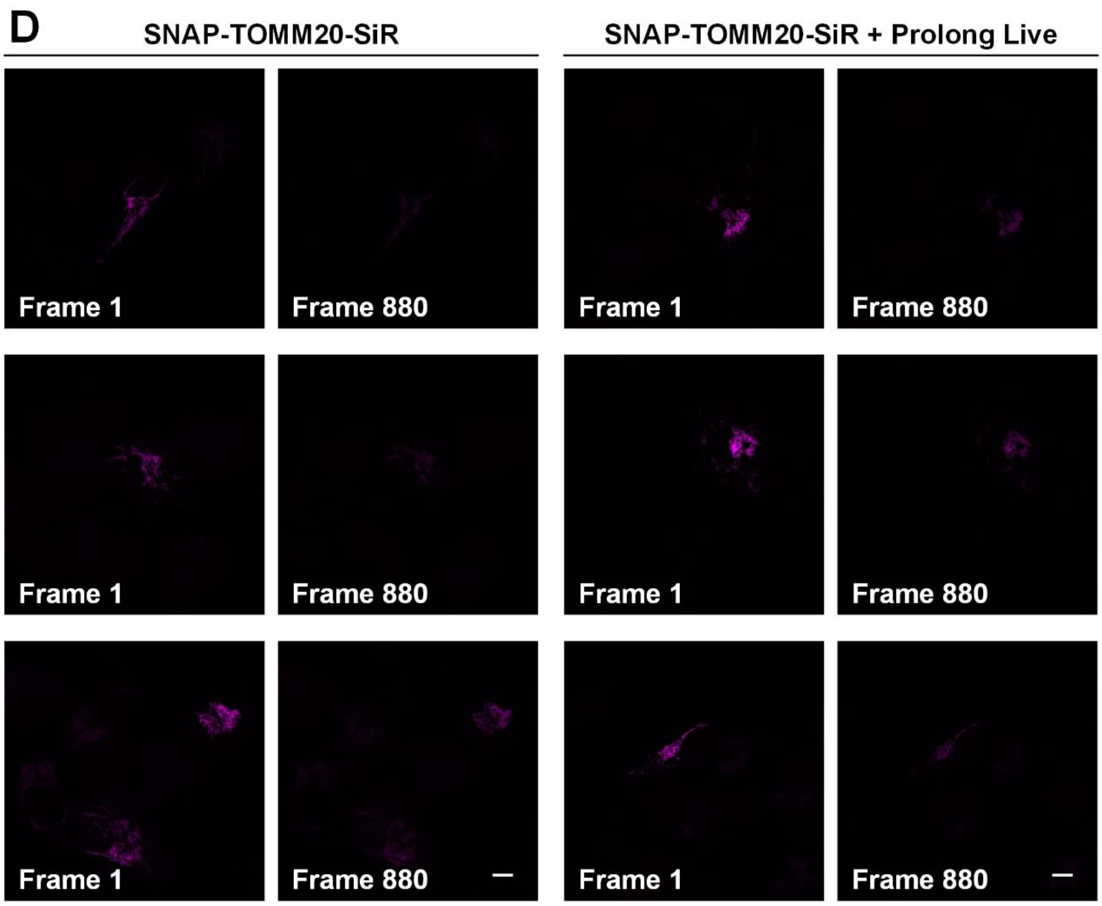
D
SNAP-TOMM20-SiR
SNAP-TOMM20-SiR + Prolong Live
Frame 1
Frame 880
Frame 1
Frame 880
Frame 1
Frame 880
Frame 1
Frame 880
Frame 1
Frame 880
Frame 1
Frame 880

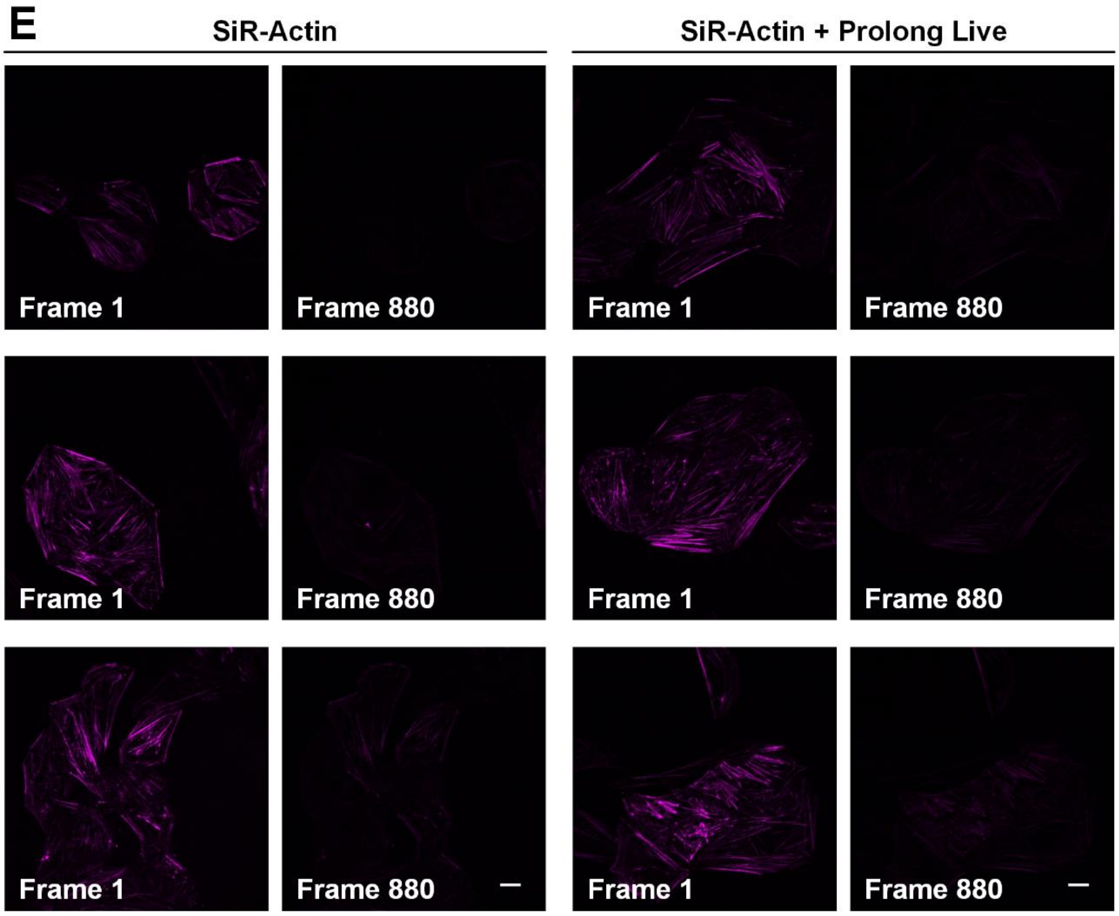


**Figure S10**. The first and last frames from 20-min confocal imaging of different probes.

Living U2OS cells were labeled with (**A**) Atto 647N, (**B**) MitoTracker Deep Red, (**C**) SNAP-Cox8A-SiR, (**D**) SNAP-TOMM20-SiR, or (**E**) SiR-Actin. Scale bars: 10 μm.

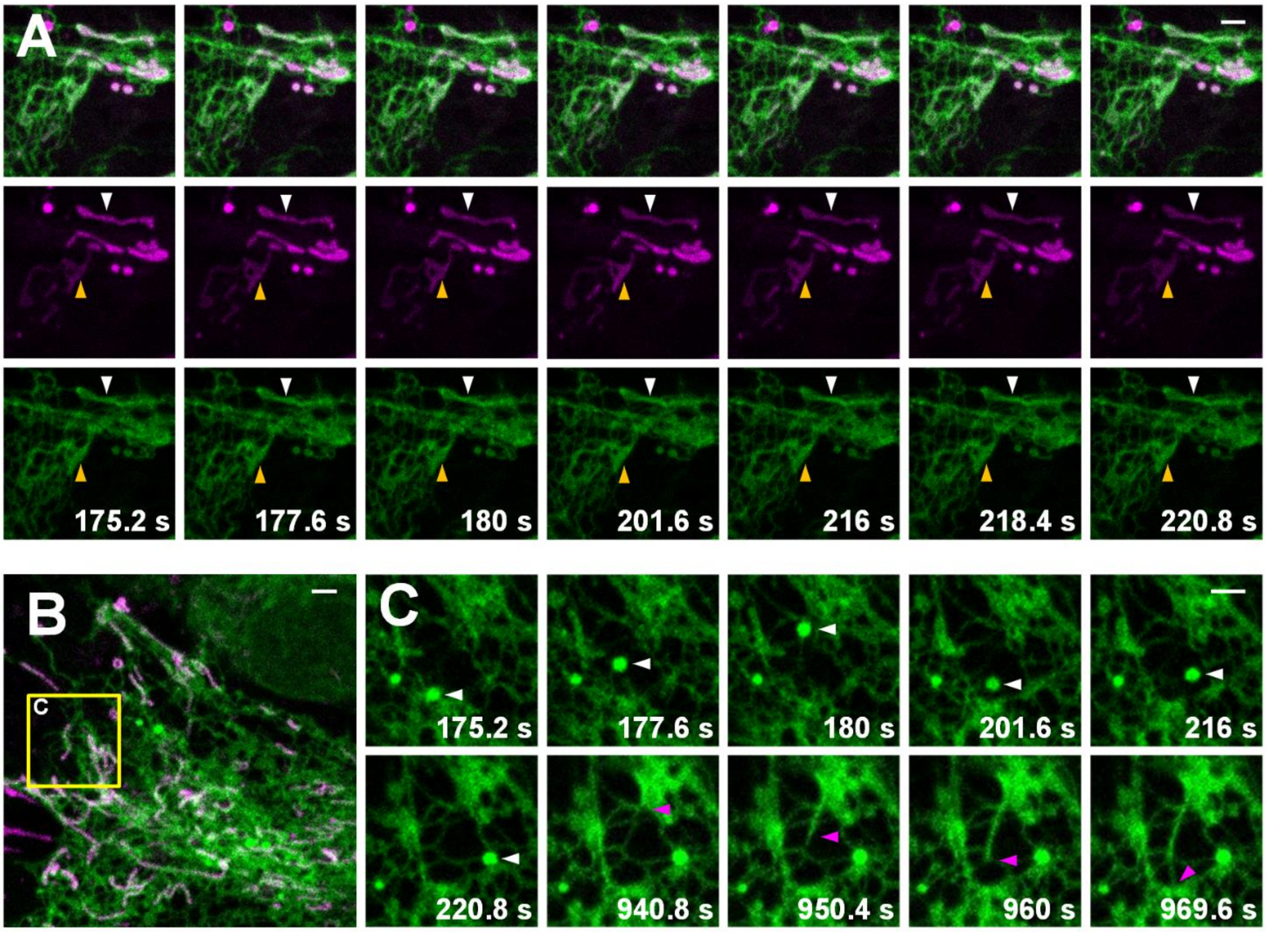


**Figure S11**. Dual-color confocal images of living Astrocytes.

(**A, B**) The cells were labeled with ER-Tracker Green (green; 2 μM) and Atto 647N (magenta; 15 μM) for 30 min at 20°C. (**C**) Vesicles or ER tubules move along ER tubules. For the time-lapse images, consecutive frames spaced at 2.4-s intervals were obtained; representative images of consecutive frames are displayed. Scale bars: 2 μm.

**Table S1**. Comparison of the optical properties of frequently-used red-absorbing fluorescent dyes.

| *Dye* | *Excitation Maximum (nm)*[a] | *Emission Maximum (nm)*[a] | *Extinction* ($m^{-1}$ $cm^{-1}$)[b] | *Quantum Yield*[c] | *Brightness*[d] |
|---|---|---|---|---|---|
| ***Atto 647N*** | 644 | 669 | 150,000 | 0.65 | 97,500 |
| ***MitoTracker Deep Red FM*** | 644 | 665 | - | - | - |
| ***SiR*** | 645 | 661 | 100,000 | 0.39 | 39,000 |
| ***BODIPY 650/665*** | 646 | 660 | 102,000 | 0.46 | 46,920 |
| ***Cy5*** | 649 | 670 | 250,000 | 0.28 | 70,000 |
| ***STAR 635P*** | 634 | 654 | 120,000 | 0.90 | 108,000 |
| ***STAR 635*** | 635 | 659 | 120,000 | 0.25 | 30,000 |
| ***Alexa Fluor 647*** | 650 | 665 | 239,000 | 0.33 | 78,870 |
| ***Atto 647*** | 645 | 669 | 120,000 | 0.20 | 24,000 |

[a] Excitation and emission peak wavelengths of dye spectra.

[b] Extinction coefficients reported by the dye manufacturers.

[c] Quantum yields from either the dye manufacturer or from the McNamara fluorophore data tables. -, values not available from the dye manufacturer or McNamara data tables.

[d] Brightness = Extinction × Quantum Yield.

**Table S2**. Experimental conditions for long-term confocal imaging.

<table>
<tr><th>Label</th><th>Dosage (μM)</th><th>Incubation temperature (°C)</th><th>Incubation time (min)</th><th>Excitation λ (nm)</th><th>Illumination intensity (kW/cm²)</th></tr>
<tr><td>Atto 647N</td><td>1.5</td><td>20</td><td rowspan="4">30</td><td rowspan="4">640</td><td rowspan="4">1.25</td></tr>
<tr><td>MitoTracker Deep Red</td><td>0.2</td><td>37</td></tr>
<tr><td>SNAP-Cell 647-SiR</td><td>3</td><td>37</td></tr>
<tr><td>Actin-SiR</td><td>1</td><td>37</td></tr>
<tr><th>Dimension</th><th>Pixel size (μm)</th><th>Exposure time per raw image (ms)</th><th>Pinhole size (μm)</th><th>Time points</th><th>Cycle time (Acquisition + resting time) (s)</th></tr>
<tr><td>512 × 512</td><td>0.25</td><td>293.92</td><td>30.00</td><td>880</td><td>1.36</td></tr>
</table>
